# Continuous flexibility analysis of SARS-CoV-2 Spike prefusion structures

**DOI:** 10.1101/2020.07.08.191072

**Authors:** Roberto Melero, Carlos Oscar S. Sorzano, Brent Foster, José-Luis Vilas, Marta Martínez, Roberto Marabini, Erney Ramírez-Aportela, Ruben Sanchez-Garcia, David Herreros, Laura del Caño, Patricia Losana, Yunior C. Fonseca-Reyna, Pablo Conesa, Daniel Wrapp, Pablo Chacon, Jason S. McLellan, Hemant D. Tagare, Jose-Maria Carazo

**Author notes:** Equally contributing authors (Roberto Melero, Carlos Oscar S. Sorzano, Brent Foster).

## Abstract

With the help of novel processing workflows and algorithms, we have obtained a better understanding of the flexibility and conformational dynamics of the SARS-CoV-2 spike in the prefusion state. We have re-analyzed previous cryo-EM data combining 3D clustering approaches with ways to explore a continuous flexibility space based on 3D Principal Component Analysis. These advanced analyses revealed a concerted motion involving the receptor-binding domain (RBD), N-terminal domain (NTD), and subdomain 1 and 2 (SD1 & SD2) around the previously characterized 1-RBD-up state, which have been modeled as elastic deformations. We show that in this dataset there are not well-defined, stable, spike conformations, but virtually a continuum of states moving in a concerted fashion. We obtained an improved resolution ensemble map with minimum bias, from which we model by flexible fitting the extremes of the change along the direction of maximal variance. Moreover, a high-resolution structure of a recently described biochemically stabilized form of the spike is shown to greatly reduce the dynamics observed for the wild-type spike. Our results provide new detailed avenues to potentially restrain the spike dynamics for structure-based drug and vaccine design and at the same time give a warning of the potential image processing classification instability of these complicated datasets, having a direct impact on the interpretability of the results.

## Introduction

The world lives in the middle of truly unexpected times, with a viral global pandemic caused by SARS-CoV-2. Science works around the clock to provide answers to essential questions aimed at understanding how viral infection occurs and how we could interfere with it. In this context, one of the most pressing issues is to analyze how the initial event of cellular recognition occurs between the viral spike (S) protein and the ACE2 receptor, aiming to start understanding the structural flexibility involved in the process. This is an essentially dynamic event, hard to analyze by most structural biology techniques. Still, cryo-EM offers some unique capabilities that makes it a very suitable approach for the task, including that it can work with non-crystalline samples and, up to a certain degree, with structural flexibility (Dashti et al., 2014; Maji et al., 2020; Scheres et al., 2007; Sorzano et al., 2019; Tagare et al., 2015).

In turn, cryo-EM information is complex, buried in thousands of very noisy movies, making it a real challenge to reveal a three-dimensional (3D) structure from this collection of images. Furthermore, cryo-EM is in the middle of a methodological and instrumental “revolution” (Kühlbrandt, 2014) that is already lasting several years, implying that new methods are being constantly produced. In a way, we can say that almost anything is “old” by the time it reaches the hands of the practitioner, and this work is a very good example of this phenomenon. In this way, the original data of Wrapp et al. (2020) have been reanalyzed applying newer workflows and algorithms, obtaining improved information.

Considering that we were studying a biological system characterized by its continuous flexibility, we have not strictly followed the standard multi-class approach (Scheres et al., 2007), very well suited to discrete flexibility cases, since the mathematical modeling and the biological reality could be just too far apart. Instead, we have calculated a new “ensemble” map at 3Å global resolution in which bias has been carefully reduced, followed by both a 3D classification process and a continuous flexibility analysis in 3D Principal Component (PC) space using a GPU-accelerated and algorithmically-improved version of the method of Tagare et al. (2015). The ensemble map has been used for atomic modeling. Our aim has been to explore a larger part of the structural flexibility present in the data set than the one achievable by 3D classification alone. Using this mixed procedure, and through the scatter plots of the projection of the different particle images onto the principal component axes, we have clearly shown how the spike flexibility in this dataset should be understood as a continuum of states rather than having discrete conformations. Thanks to maximum likelihood-based classification we have obtained two maps that project at the extremes of the main principal component on which flexible fitting from the ensemble map has been performed. Still, these extreme maps have an intrinsic blurring on the most flexible areas, since for any class we may define, images are coming from a continuum of states and are, therefore, heterogeneous. This flexibility is substantially reduced in a recently described biochemically stabilized spike (Hsieh et al., 2020), as evidenced by the reduced blurring that translates into an improved local resolution.

In this work, we describe the new structural information obtained and how it impacts our biological understanding of the system, together with the new workflows and algorithms that have made this accomplishment possible. At the same time, we are currently submitting our raw and intermediate data, including preprocessing workflows, to public databases (EMPIAR (Iudin et al., 2016) and EMDB (Lawson et al., 2011)) with the hope to further speed up developments and to enhance scientific reproducibility.

## Results

With the goal set at analyzing spike flexibility, we go step by step over our key results.

### Ensemble map and the way to obtain it

In the following, we describe the analysis of the spike stabilized in the prefusion state by two proline substitutions in S2 (S-2P) or a more recent variant containing six proline substitutions in S2 (HexaPro). We will objectively demonstrate that the spike’s flexibility should be understood as a quasi-continuum of conformations, so that when performing a structural analysis on this specimen special care has to be paid to the images processing workflows, since they may directly impact the interpretability of the results.

Starting from the original SARS-CoV-2 S-2P data set of Wrapp et al. (2020), we have completely reanalyzed the data in the context of our public domain software integration platform Scipion (de la Rosa-Trevín et al., 2016), breaking the global 3A resolution barrier. A representative view of the new ensemble map and its corresponding global FSC curve is shown in Figure 1A (new EMD-11328); the sequence of a monomer of the S protein is shown on the right to facilitate further discussions on structure-function relationships (from Wrapp et al. (2020)). Figures 1B and 1C show a comparison between the original map (Wrapp et al., 2020) with EMDB entry 21375 and the newly reconstructed ensemble map corresponding to EMD-11328. Clearly, local resolution (Vilas et al., 2018) -left- is increased in the new map, and anisotropy-center-is much reduced. Finally, on the right-hand side, we present plots of the radially-averaged tangential resolution, that are related to the quality of the angular alignment (Vilas et al., 2020); the steeper the slope, the higher the angular assignment error. As can be appreciated, the slope calculated from the newly obtained map is almost zero, compared with Wrapp et al. (2020), indicating that, in relative terms, the particle alignment used to create the new map is better than the one used to build the original map. The result is an overall quantitative enhancement in map quality.

**Figure 1.**
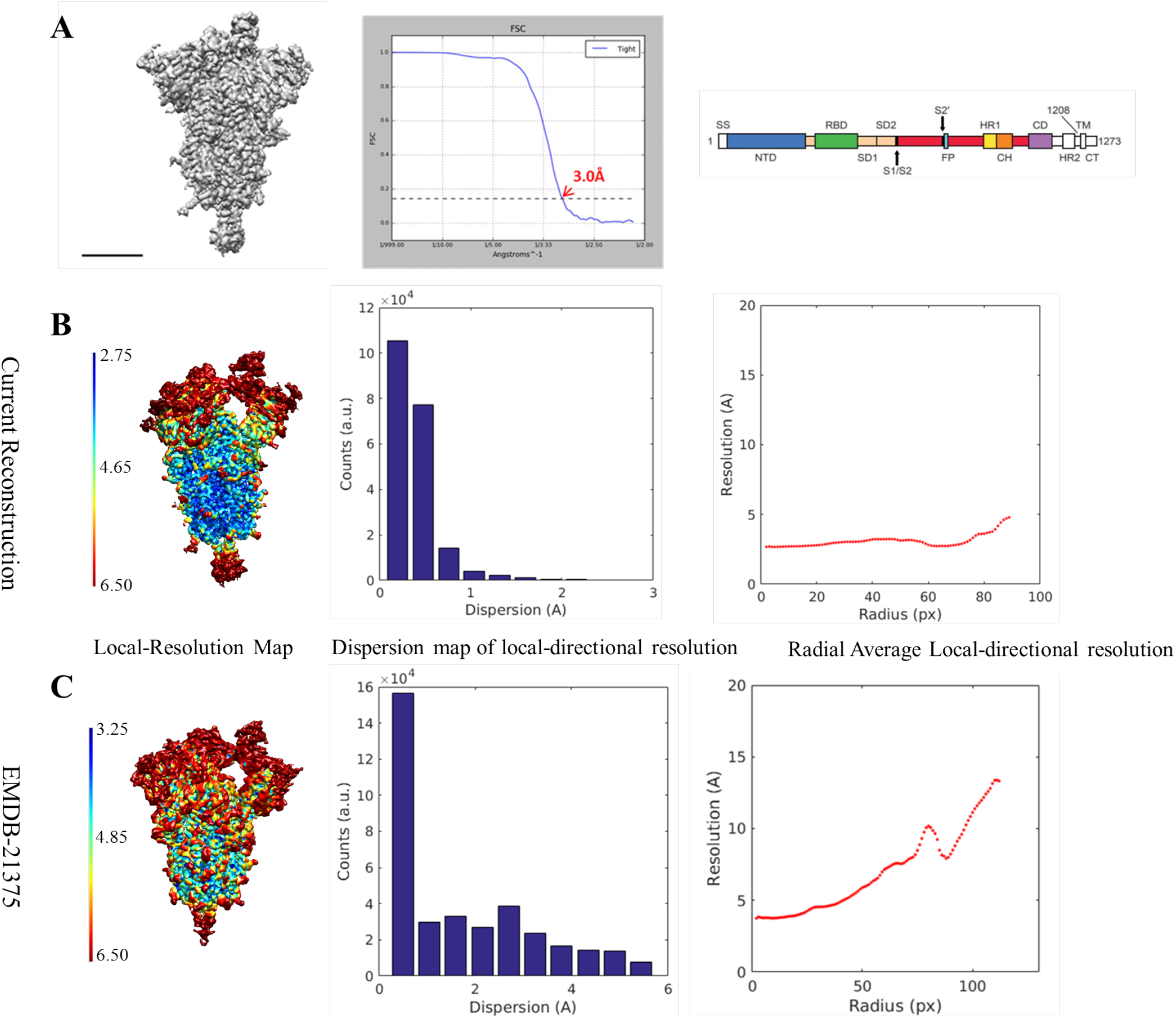
The spike and the ensemble map. A) A representative view of the new map (EMD-11328), the corresponding FSC curve and the sequence of a monomer of the S protein (from Warpp et al., (2020)). Scale bar 5 nm. B-C) New ensemble cryo-EM map (EMD-11328) compared with the one originally presented (EMDB 21375). The first line (B) corresponds to the new map and the second one (C) to EMD-21375. Within each line, and from left to right: Map representation showing local resolution, histogram representation of local directional resolution dispersion (interquartile range between percentiles 17 - 83) and, finally, plot showing radial average of local tangential resolution.

In terms of tracing, besides modeling several additional residue side chains and improving the geometry of the carbon skeleton (see Supplementary Material Figure SM2), one of the most noticeable improvements that we observed in the new map is the extension of the glycan chains that were initially built, particularly throughout the S2 fusion subunit (new PDB 6ZOW). A quantitative comparison can be made between the length of glycan chains in the new “ensemble structure” with respect to the former one (PDBID: 6VSB) (see Supplementary Table SM2). Although the total number of *N*-linked glycosylation sequons throughout the SARS-CoV-2 S trimer is essentially the same in the new structure (45) and in 6VSB (44), we have substantially increased the length of their glycan chains, expanding the total number of glycans by about 50%. We note the importance of this extensive glycosylation for epitope accessibility, and how the accurate determination of this glycan shield will facilitate efforts to rapidly develop effective vaccines and therapeutics. Supplementary Material Figure SM2 shows a representative section of sharpened versions of ensemble map (EMD-11328) as compared to EMD-21375 where glycans can be better traced now. Still, we should not forget that the ensemble map contains images in which the receptor-binding domain (RBD) and N-terminal domain (NTD) are in different positions (see next section), and consequently, these domains appear blurred. Details on how the tracing was done can be found in Materials and Methods, while in Supplementary Material Figure SM3 we present two maps-to-model quality figures indicating the good fit, in general, with the obvious exception of the variable parts.

### Flexibility analysis

Starting from a carefully selected set of particles obtained from our consensus and cleaning approaches (see Material and Methods), together with the ensemble map described previously, we subjected the data to the following flexibility analysis:

1. The original images that were part of the ensemble map went through a “consensus classification” procedure aimed at separating them into two algorithmically stable classes. Essentially, and as described in more detail in Material and Methods, we performed two independent classifications, further selecting those particles that were consistently together through the two classifications. In this way, we obtained two new classes shown in Figure 2A. We will refer to them as “the closed conformation” (Figure 2A-Class1; EMD-11336) and “the open conformation” (Figure 2A-Class2; EMD-11337). The number of images in each class was reduced to 45k in one case, and 21k in the other, with global FSC-based resolutions of 3.1 and 3.3 Å, respectively. The open and closed structures depict a clear and concerted movement of the “thumb” formed by receptor-binding domain (RBD) and subdomain 1 and 2 (SD1 & SD2) and the NTD of an adjacent chain. The thumb moves away from the central spike axis, exposing the RBD in the up conformation. In order to make clearer where the changes are at the level of Class 1 and Class 2 maps, we have made use of Sorzano et al. (2016) representation of map local strains, that help visualize very clearly the type of strains needed to relate two maps, whether it is rigid body rotations or some more complex deformations are needed (stretching). We have termed the maps resulting from this elastic analysis as ‘1s’ (Class 1, stretching) and ‘1r’ (Class 1, rotations) on the right hand side of Figure 2A, and the same for Class 2. The color scale in both stretching and rotations goes from blue (small) to red (large). Clearly the differences among the classes with respect to the NTD and RBD have a very substantial component of pure coordinated rigid body rotations, while the different RBDs present a much more complex pattern of deformations (stretching), indicating an important structural rearrangement in this area that does not happen elsewhere in the specimen. In terms of atomic modeling, we have made a flexible fitting of the ensemble model onto the closed and open forms (see Figure 2A, rightmost map; the PDB ID for the open conformation is PDB ID 6ZP7, while for the closed one it is PDB ID 6ZP5). Focusing on rotations, which is the most simple element to follow, we can quantify that the degree of rotation of the thumb in these classes is close to 6 degrees, as shown in Figure 2B. Given this flexibility, we consider that the best way to correctly present the experimental results is through the movie shown in Supplementary Material Video 1, where maps and atomic models are presented. Within the approximation to modeling that a flexible fitting represents, we can appreciate two hinge movements at RBD-SD1-2 domains: one located between amino acids 318 to 326 and 588 to 595 that produces most of the displacement, and other between amino acids 330 to 335 and 527 to 531 that goes together with a less pronounced “up” movement of the RBD. This thumb motion is completed by the accompanying motion of the NTD from an adjacent chain. Also in a collective way, other NTDs and down RBDs are slightly moving, as can be appreciated better in the S1 movie where the transition between fitted models overlaps with the interpolation between observed high-resolution class maps.
2. To further investigate whether or not the flexibility was continuous, we proceeded as follows: Images from the two classes were pooled together and, using the ensemble map, subjected to a 3D principal components analysis (PCA). The approach we followed is based on Tagare et al. (2015), with some minor modifications of the method. A detailed explanation of the modifications is given in Material and Methods. We initialized the first principal component to the difference in the open and closed conformation, while the remaining principal components were initialized randomly. Upon convergence, the eigenvalue of each principal component and the scatter of the images in the principal component space was calculated. The eigenvalues of the principal components are shown in Figure 3A. Clearly the first three principal components are significant. The scatter plot of the image data in Principal Component 1-3 space is shown in Figure 3B. Figure 3B strongly suggests that there is “continuous flexibility” rather than “tightly clustered” flexibility. Figure 3B also shows the projection of the maps corresponding to the open and closed conformations on the extremes of the first three Principal Components. It is clear that the open and closed conformations are aligned mostly along the first Principal Component; suggesting that the open/close classification captures the most significant changes. Figure 3C shows side views of a pair of structures (mean plus/minus 2 x std, where std=sqrt(eigenvalue)) for each Principal Component. Additional details of these structures are available in the Supplementary Material Figures SM4 and SM5. Note that Principal Components are not to be understood as structural pathways with a biological meaning, but directions that summarize the variance of a data set. For instance, the fact that RBD appears and disappears at the two extremes of PC3 indicates that there is an important variability in these voxels, probably indicative of the up and down conformations of the RBD (to be understood in the context of the elastic analysis shown in Figure 2B).
3. Through this combination of approaches, we have learnt that the spike conformation fluctuates virtually randomly in a rather continuous manner. Additionally, clearly the approach taken to define the two algorithmically stable “classes” has partitioned the data set according to the main axis of variance, PC1, since the projection of these classes’ maps fall almost exclusively along PC1 and are located towards the extremes of the image projection cloud. Note that the fraction of structural flexibility due to PC2 and PC3 is also important in terms of the total variance of the complete image set, but that classification approaches do not seem to properly explore it. Unfortunately, currently the resolution in PC2 and PC3 is limited, so it is difficult to derive clear structural conclusions from these low resolution maps. Still, it is clear from this data that the dynamics of the spike is far richer than just a rigid body closing and opening, and involves more profound rearrangements, especially at the RBD but at other sites as well. This observation is similar to the one of Ke et al. (2020), working with subtomogram averaging.

**Figure 2.**
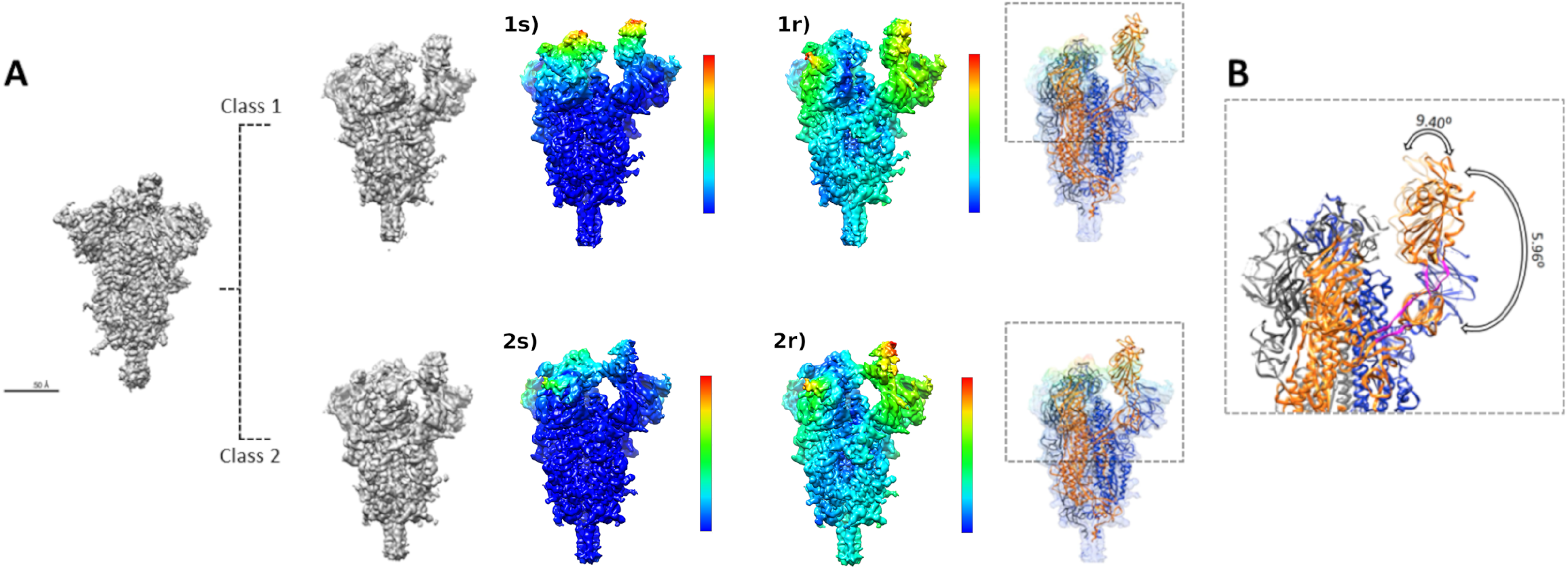
Flexibility analysis: A) A representative view of the new ensemble map and the two new classes showing in Class 1 “the open conformation” and in Class 2 “the closed conformation”. Note the elastic analysis of deformations on the Class 1 and Class 2 maps (see main text), with 1s) referring to “stretching” and 1r) to “rotations”. Color code goes from blue (minimal deformation) to red (maximal deformation). B) Representation of the angles defined by the spike when transitioning between the opened and the closed states. The regions shown in magenta represent the hinges used by the RBD domain to pivot. The first hinge spans amino acids 318 to 326 and 588 to 595, while the second hinge is defined by aminoacids 330 to 335 and 527 to 531. The angles were measured using PyMol software.

**Figure 3.**
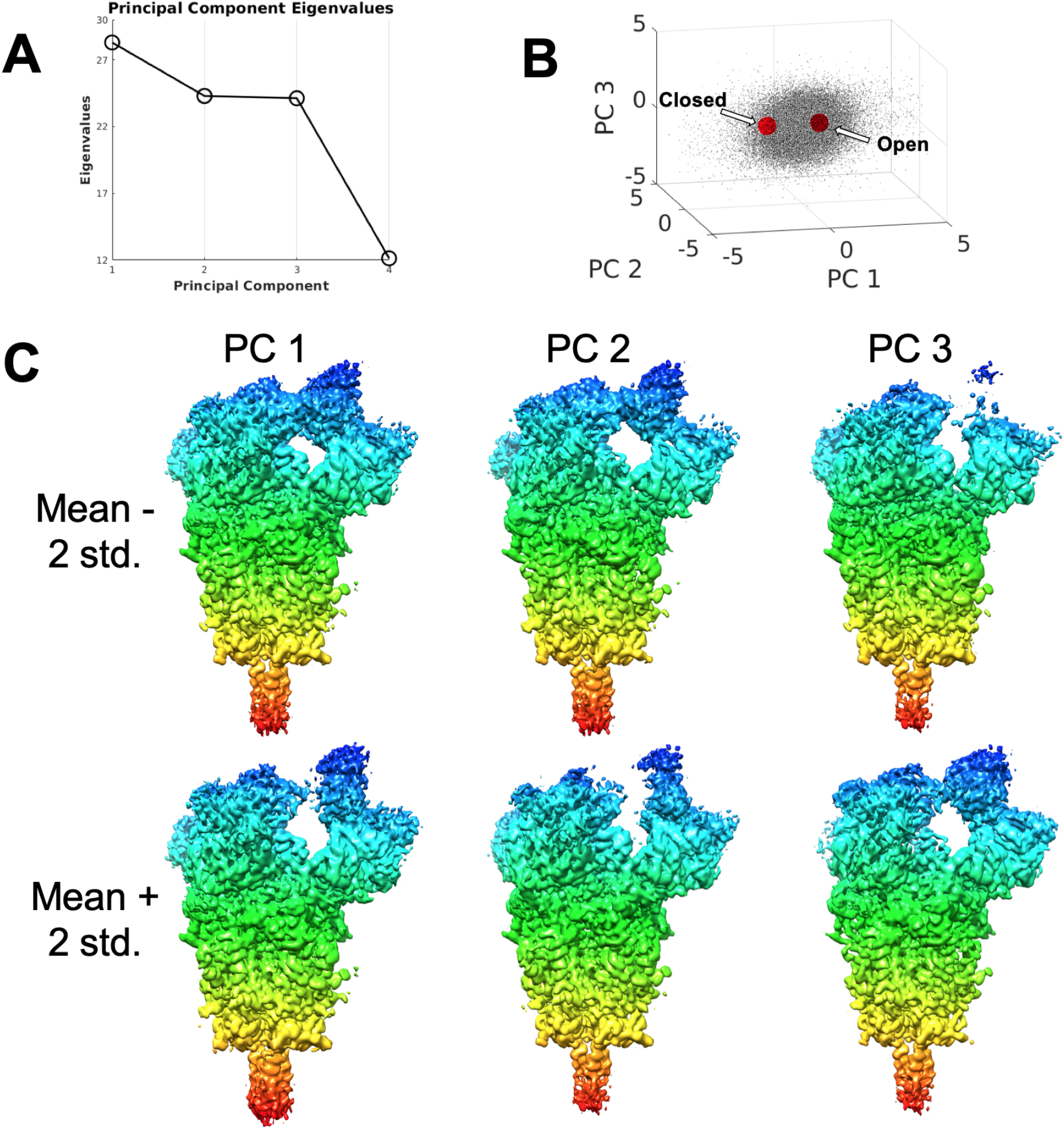
Principal Component Analysis of the Cov-2 spike structure. A) Eigenvalues of principal components. The first three principal components are significant. B) Scatter plot of the contribution of the first three principal components to each particle image together with the projection of the open and closed class maps, shown as red points. The difference between the projections of the two maps is mostly aligned along PC1. C) Side view of the first two principal components shown as mean +/−2 times std, where std=sqrt(eigenvalue). Coloring indicates z-depth of the structure, and is added to assist visualization. Supplementary Material Figures 4 and 5 contain additional views of these structures.

Additionally, the fact that PCA indicates this continuous flexibility as a key characteristic of the spike dynamics also suggests that many other forms of partitioning (rather than properly “classifying”) this continuous data set could be devised, this fact just being a consequence of the intrinsic instability created by forcing a quasi-continuous data distribution without any clustering structure to fit into a defined set of clusters. In this work we have clearly forced the classification to go to the extremes of the data distribution -as shown in Fig. 3-, probably by enforcing an algorithmic stable classification, but the key result is that any other degree of movement of the spike in between these extremes of PC1 as well as PC2 and PC3 would also be consistent with the experimental data. In other words, since the continuum of conformations does not have clear “cutting/classification” points, there is a certain algorithmic uncertainty and instability as to the possible results of a classification process. Note that this instability could be exacerbated by the step of particle picking, in the sense that different picking algorithms may have different biases (precisely to minimize this instability we have done all throughout this work a “consensus” approach to picking).

Clearly, flexibility is key in this system, so that alterations in its dynamics may cause profound effects, including viral neutralization, and this could be one of the reasons for the neutralization mechanism of antibodies directed against the NTD (Chi et al., 2020).

### Structure of a biochemically stabilized form of the spike

In this work we have also analyzed the HexaPro stabilized spike in the prefusion state (Hsieh et al., 2020). In this case, and after going through the same stringent particle selection process than for the previous specimen, which is presented in depth in Material and Methods, it was impossible to obtain stable classes, so that in Fig. 4 we present a single map (EMD-11341), together with its global FSC curve and a local resolution analysis. It is clear that local resolution has increased in the moving parts (mostly RBD and NTD), although we did not still feel confident for further modeling.

**Figure 4.**
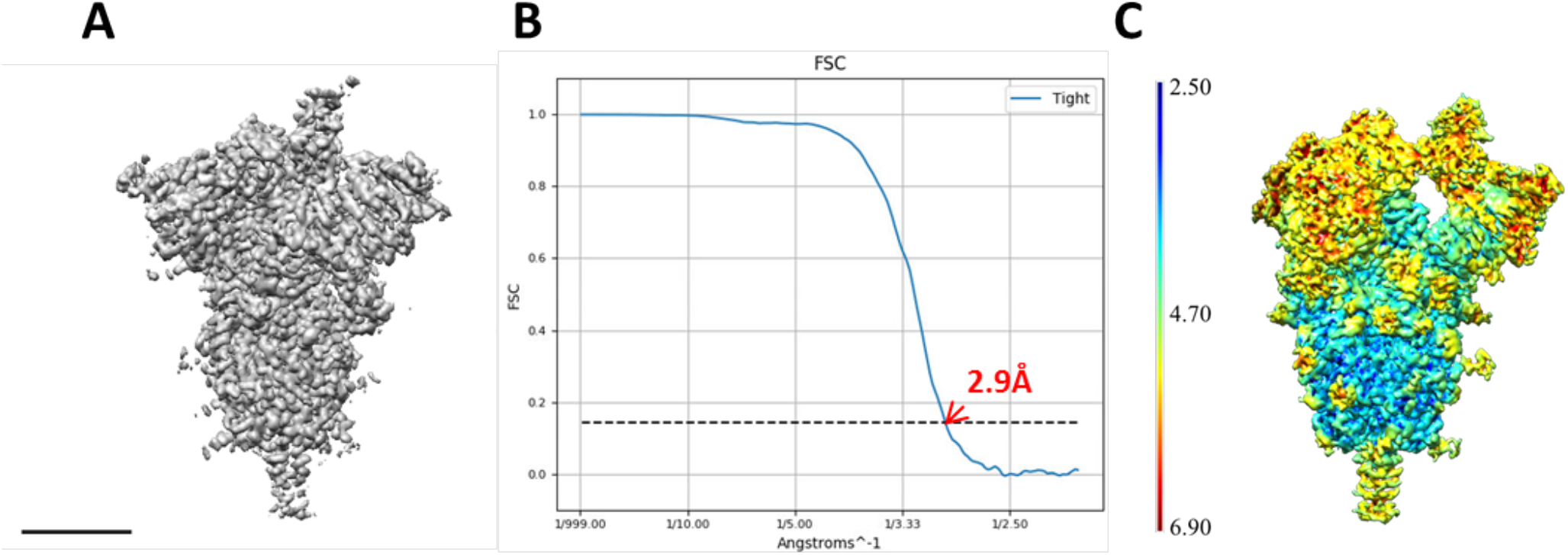
Analysis of a biochemically stabilized form of the spike. A-B) A representative view of the stabilized form of the spike map and the corresponding FSC curve. Scale bar 5 nm. C) Local resolution map estimated with MonoRes.

## Conclusions

We present in this work a clear example of how the structural discovery process can be greatly accelerated by a wise combination of fast data sharing and the use of the wave of newly developed algorithms that characterize this phase of the “cryo-EM revolution”. The reanalysis of the same data used in Wrapp et al. (2020), but with new workflows and new tools, has resulted in a rich analysis of the spike flexibility as a key characteristic of the system.

Essentially, and at least to a first approximation, the spike moves in a continuous manner with no preferential states, as clearly shown in the scatter plots of Figure 3B. In this way, the results of a particular instance of image processing analysis, including a 3D classification, should be regarded as snapshots of this quasi-continuum of states. In our case we have shown that a particular meta image classification approach, implemented through a consensus among different methods in many steps of the analysis, results in classes that are at the extreme of the main axis of variance in Principal Component space. Clearly PC1, through the analysis of the two extreme classes, reflects a concerted motion of the NTD-RBD-SD1-2 thumb, although there are smaller collective movements all throughout the spike (see Fig. 2 and Supplementary Material Video 1). In this case, the RBD moves together with the NTD, with a smaller degree of independent flexibility and always in the “up” conformation. The NTD-RBD movement can be characterized to a large degree as a rotation, but the different RBDs present a much more complex pattern of flexibility, indicating an important structural rearrangement (from Figure 2, elastic analysis, and Figure 3, PCA). The presence of quasi-solid body rotation hinges is clearly located between amino acids 318 to 326 and 588 to 595, that produces most of the displacement, together with other hinges between amino acids 330 to 335 and 527 to 531, that goes together with a less pronounced “up” movement of RBD

Still, there are other Principal Component axes explaining significant fractions of the inter-image variance that are not properly explored at the level of our two classes. Principal Component 3 is a clear example, indicating a high variance at the voxels associated with RBD up, which is probably suggesting large conformational changes in that area that result in RBD coming down.

The flexibility analysis performed in this work complements previous analysis showing large rotations together with RBD up-down structural changes (Pinto et al., 2020; Wrapp et al., 2020), in the sense that the different studies present “snapshots” of a continuum of movements obtained by a particular instance of an image processing classification. In a sense, all these results are correct, but none of them is able to capture the flexibility richness of this system. This fact reflects the intrinsic instability of segmenting a continuum into defined clusters, which is a clear limitation of classification approaches to be considered in the detailed analysis of any dataset from this system.

An obvious way to increase resolution in the moving parts of the spike is to reduce their mobility, which is the case, for instance, of the biochemical stabilization of Hsieh et al. (2020), and also of the formation of a complex with an antibody against NTD (Chi et al., 2020). On the other hand, the way towards a more complete analysis of the flexibility of the spike necessarily involves the analysis of quite substantially larger datasets than those being used in most current CoV-2 studies, so that all the main axes of inter-image variability can be explored, which is work under development at the moment.

From a biomedical perspective, the proof that a quasi-continuum of flexibility is a key characteristic of this specimen, a concept implicitly considered in much of the structural work performed so far but never demonstrated, suggests that ways to interfere with this flexibility could be important components of new therapies.

## Acknowledgements

We acknowledge the support from the Advanced Computing and e-Science group at the Institute of Physics of Cantabria (IFCA-CSIC-UC) as well as the Barcelona Supercomputer Center (access project BCV-2020-2-0005). The authors would like to acknowledge financial support from: CSIC, (PIE/COVID-19 number 202020E079), the Comunidad de Madrid through grant CAM (S2017/BMD-3817), the Spanish Ministry of Science and Innovation through projects (SEV 2017-0712, FPU-2015/264, PID2019-104757RB-I00 (AEI/FEDER, UE), BFU2016-76220-P and PID2019-109041GB-C21 (AEI/FEDER, UE), the Instituto de Salud, Carlos III, PT17/0009/0010 (ISCIII-SGEFI / ERDF) and the European Union and Horizon 2020 through grant: INSTRUCT - ULTRA (INFRADEV-03-2016-2017, Proposal: 731005), EOSC Life (INFRAEOSC-04-2018, Proposal: 824087), HighResCells (ERC - 2018 -SyG, Proposal: 810057), IMpaCT (WIDESPREAD-03-2018 - Proposal: 857203) and EOSC – Synergy (EINFRA-EOSC-5, Proposal: 857647) The authors HDT and BF were supported by the NIH Grant GM125769 and JSM was supported by NIH grant R01-AI127521. The authors acknowledge the support and the use of resources of Instruct, a Landmark ESFRI project

## Authors Contribution

RME (Roberto Melero) and COSS have performed all the image analysis in Scipion, while BF has done an equivalent work for Principal Component Analysis and JLV the local resolution analysis. MM and RMA (Roberto Marabini) have been in charge of structural modelling, while PCh (Pablo Chacon) has performed the flexible fittings and incorporated important sections of the manuscript. ER-A performed the map-to-model analysis as well as sharpened cryo-EM maps, while RS-G also worked in new sharpening methods and DH performed the elastic inter class analysis. PCO (Pablo Conesa), YF-R, LdC and PL have been in charge of the IT hardware and software support. JMcL and DW have supplied the images and provided advice throughout the work. HT, COSS and JMC have conceptualized the work, with JMC making the manuscript writing that was complemented by all other authors. JMcL, HT and JMC were responsible for the funding.

## Competing interests

None

## Materials and Methods

### Image Processing Workflow

The basic elements of the workflow combine quite classic cryo-EM algorithms with recent improvements in particle picking (Sanchez-Garcia et al., 2020b, 2018; Wagner et al., 2019) and key ideas of meta classifiers, which integrate multiple classifiers by a “consensus” approach (Sorzano et al., 2000), finalizing with a totally new approach to map post-processing based on deep learning that we term “Deep cryo EM Map Enhancer” (Sanchez-Garcia et al., 2020a), that complements our previous proposal on local deblurring (Ramírez-Aportela et al., 2020b). Naturally, map and map-model quality analysis are performed with a variety of tools (Pintilie et al., 2020; Ramírez-Aportela et al., 2020a; Vilas et al., 2020). Conformational variability analysis is carried out explicitly addressing the continuous flexibility nature of the underlying biological reality, in which SARS-CoV-2 spike is exploring the conformational space to bind the cellular receptor. Most of the image processing done in this work has been done using Scipion framework (de la Rosa-Trevín et al., 2016) which is a public domain image processing framework freely available at url http://scipion.i2pc.es.

A graphical representation of the image processing workflow used in this work can be found in Suppl. Material Figure 1

### Meta Classifiers

On meta classifiers, and as discussed in Sorzano et al. (2020), the rationale is that a careful analysis of the ratio between algorithmic degrees of freedom versus data size shows that cryo-EM may has transitioned from an area characterized by parameter variance to one dominated by possible parameter biases. In very simple terms, we have a lot of data, so we can fight the variance in our data if we deal with random errors. However, whenever there is the possibility of a systematic error, a so-called “bias”, artifacts in the maps may occur and, if this is the case, they can be very difficult to detect. We deal with the problem of introducing bias in the map through “consensus”, so that we select those parameters for which several methods, as methodologically “orthogonal” as possible, concur on the same answer (sometimes we also use different runs of the same method).

This notion has been used at several different steps of the workflow. In particular:

1. CTF estimation: We estimated the microscope defocus using two different programs (GCTF (Zhang, 2016) and CTFFind4 (Rohou and Grigorieff, 2015). We only selected those micrographs for which both estimates agreed up to 2.1 Å (Marabini et al., 2015).
2. Particle selection: We used two particle picking algorithms (Xmipp (Abrishami et al., 2013) and Cryolo (Wagner et al., 2019)). We submitted both results to a picking consensus algorithm by deep learning (Sanchez-Garcia et al., 2018) and removed all those coordinates in contaminations, carbon edges,… also using a deep learning algorithm (Sanchez-Garcia et al., 2020b). Then we cleaned the set of selected particles using two rounds of CryoSparc 2D classification (Punjani et al., 2017; Punjani and Fleet, 2020) and the consensus of two independent 3D classifications with CryoSparc.
3. Initial volume: As initial volume we selected the majoritarian class that came out from the two 3D classifications above and refined it with Highres (Sorzano et al., 2018) with a local refinement of the 3D alignment.
4. 3D reconstruction: We then performed a CryoSparc non-uniform 3D reconstruction, followed by a local angular refinement using Relion with a 3D mask (Zivanov et al., 2018). Particle images were subjected to ctf refinement and Bayesian polishing (Zivanov et al., 2018), before performing another two rounds of ctf refinement and local angular refinement in Relion, where we improved the resolution versus the first local refinement. Finally we performed a non-uniform refinement in cryoSPARC. The reported nominal resolution 2.96Å is based on the gold-standard Fourier shell correlation (FSC) of 0.143 criterion. Actually, by using Xmipp Highres (Sorzano et al., 2018) we could lower the resolution to 2.2Å in the central region of the volume (the one that is not flexible), but at the expense of still reducing it more in the flexible areas.
5. 3D classification: We then performed two rounds of 3D classification with Relion followed by a consensus 3D class yielding two stables, large classes. With these two classes we then performed a local angular refinement using a CryoSparc non-uniform 3D reconstruction.

### Particle selection

We found that micrographs and particles that are used for the 3D reconstruction play a key role in the quality and characteristics of the final map. In particular we used the following two procedures:

a. CTF estimation: We estimated the microscope defocus using GCTF and CTFFind4. We required that both estimates are similar (the phase of their corresponding Contrast Transfer Function differed in less than 90 degrees) up to 2.1 Å. Only 70% of the micrographs met this criterion. We then estimated the CTF envelope using Xmipp CTF (Sorzano et al., 2007) while keeping fixed the defocus value (calculated as the average between the GCTF and CTFFind4 estimates). We found this step very important to keep high resolution information. With Xmipp CTF we discovered that most of the micrographs had a non-astigmatic validity between 3-4 Å (meaning that at this resolution the assumption of non-astigmatism breaked down for most of the micrographs, and only a minority of 30% reached higher resolution in a non-astigmatic way).
b. Particle selection: Two advanced particle picking algorithms were employed: Xmipp and Cryolo. The first one identified 1.2 Million (M) coordinates possibly pointing to spike particles, while the second one identified 0.73M. We then combined both estimates using Deep Consensus with a threshold of 0.99, resulting in 0.62M coordinates. Micrograph Cleaner was used to rule out particles selected in the carbon edges, aggregations or contaminations, rejecting a total amount of 50k particles. After two rounds of CryoSparc 2D classification at a pixel size of 2.1 Å and an image size of 140×140 pixels, we kept 298k particles assigned to 2D classes whose centroid clearly corresponded to projections of the spike. At this point we performed two initial volume estimates using CryoSparc and classifying the input particles into two classes. In both executions, one of the structures clearly corresponded to the spike (with 80% of particles), while the other one resulted in a 3D structure that clearly corresponded to contamination. We calculated the consensus of the two CryoSparc 3D classifications (those particles that consistently were assigned to the same 3D class). Only 203k particles belonged to the class consistently assigned to the spike.

### Validation and quality analysis

On judging the quality of our structural results, we concentrated here in three of the newest approaches: Directional Local Resolution, Q-score and FSC-Q. The first one provides a measure of map quality, while the two latter ones focus on the relationship between map and structural model. In other words, how well the model is supported by the map density, without any other complementary piece of information.

In terms of map-to-model validation, in Figure SM3A and SM3B we present Q-score and FSC-Q metrics, respectively, showing the agreement between the ensemble cryo-EM map and the structural model derived from it. In most areas the agreement is very good, with the exception of the receptor binding domain (RBD) and substantial parts of the N-terminal domain (NTD), as expected by their higher flexibility.

### Volume post-processing

In this work we have used two types of volume post-processing approaches, in the two cases they depart substantially from the traditional approach in the field that is the application of global B-sharpening. One of the approaches is our already introduced LocalDeblur sharpening method (Ramírez-Aportela et al., 2020b). The second approach is a totally new method based on deep learning (Sanchez-Garcia et al., 2020a). Concentrating on the latter method, DeepEMhancer, it relies on a common approach in modern pattern recognition, where a Convolutional Neural Network (CNN) is trained on a known data set, comprised of pairs of data points and targets, with the aim of predicting the targets for new unseen data points. In this case, the training has been done presenting the CNN with pairs of cryo-EM maps collected from EMDB and maps derived from the structural models associated with the experimental maps. As a result, our CNN learned how to obtain much cleaner and detailed versions of the experimental cryo-EM maps, improving their interpretability.

Trying to take advantage of their complementary information, we have used the two post-processed maps to trace the atomic model (PDB 6ZOW). Some examples of the similar improvement of the structure modeling according to these two sharpened maps are shown in Suppl. Mat. Figure SM2. Sharpened and unsharpened maps are all being deposited at EMDB.

### Model building

The atomic interpretation of the SARS-Cov-2 spike 3D map (PDB 6ZOW) was performed taking advantage of the modeling tools integrated in Scipion as described in Martínez et al. (2020). Due to the lack of sufficient density of the “up” conformation of the RDB, we fitted rigidly the structure of the chain A (residues 336-525) of the SARS-Cov-2 RDB in complex with CR30022 Fab (PDB ID 6YLA) to the 3D map using UCSF Chimera (Pettersen et al., 2004). This unmodeled part of the structure was called chain “a” since it was part of the chain A in the structure already inferred from the same data set (PDB ID 6VSB). The rest of the molecule was modeled using as template the same original structure (PDB ID 6VSB), as well as another spike ectodomain structure in its open state (PDB ID 6VYB). The former structure (PDB ID 6VSB) was fitted to the new map and refined using Coot (Emsley et al., 2010) and Phenix *real space refine* (Afonine et al., 2018). Validation metrics were computed to assess the geometry of the new hybrid model and its correlation with the map using Phenix *comprehensive validation (cryo-EM)*, EMRinger algorithm (Barad et al., 2015), Q-score (Pintilie et al., 2020) and FSC-Q (Ramírez-Aportela et al., 2020a). Score values considering the whole hybrid spike and excluding the unmodeled RBD are detailed in Suppl. Table SM1. The hybrid atomic structure is being submitted to EMDB.

iMODFIT (Lopéz-Blanco and Chacón, 2013) was employed to flexibly fit the hybrid atomic structure into the open and closed class maps.

### Principal component analysis

The principal component analysis follows the EM-algorithm presented in Tagare et al. (2015) with the following minor modifications: first, in contrast to Tagare et al. (2015), the images were not Wiener filtered, nor was the projected mean subtracted from the images; instead the CTF of each image was incorporated in the projection operator of that image and a variable contrast was allowed for the mean volume in each image. The extent of the variable contrast was determined by the Principal Component EM-algorithm. Second, the mean volume was projected along each projection direction and an image mask constructed with a liberal soft margin to allow for heterogeneity. The different masks thus created -one mask per projection direction-were applied to the images and the masked images were used as data. This step corresponds to imposing a form of sparsity on the data, which is known to improve the estimate of principal components in high dimensional spaces (Johnstone and Paul, 2018). All images were downsampled by a factor of 2 to improve signal to noise ratio and speed up processing. Finally, during each EM-iteration, the principal components were low pass filtered with a very broad filter whose pass band extended to 4 A. This helped in the convergence of the algorithm without significantly limiting the principal component resolution.

As part of the EM-iteration, the algorithm in Tagare et al. (2015), conveniently estimates the expected amount by which each principal component is present in each image (this is the term E[z_j] in equation 15 of Tagare et al., (2015). Figure 3B is a scatter plot of E[z_j].

---

**Supplementary Material Figure 1.**
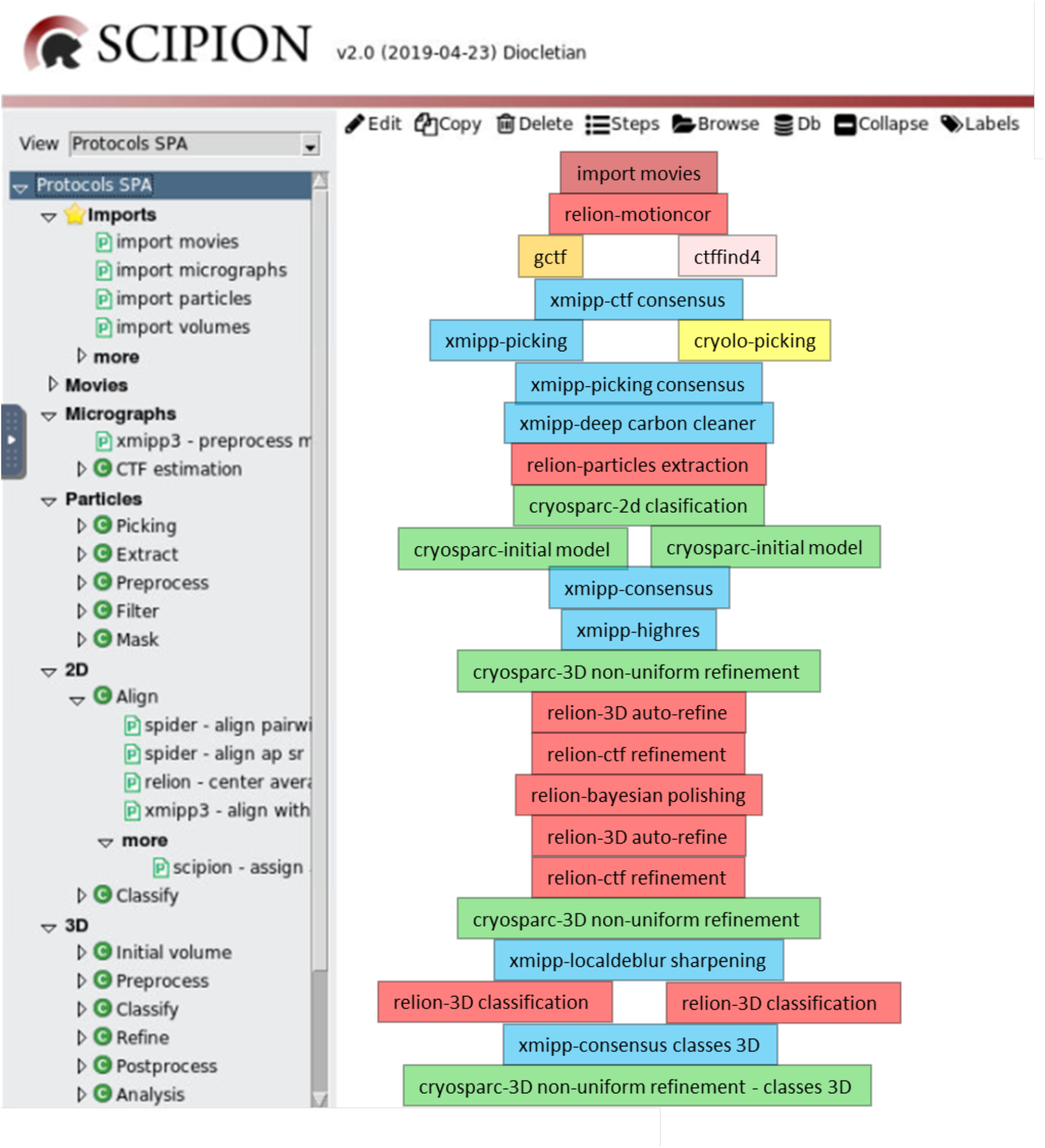
Graphical representation of the processing workflow in Scipion. The workflow is also accessible at Scipion Workflow Repository at http://workflows.scipion.i2pc.es/.

**Supplementary Material Figure SM2.**
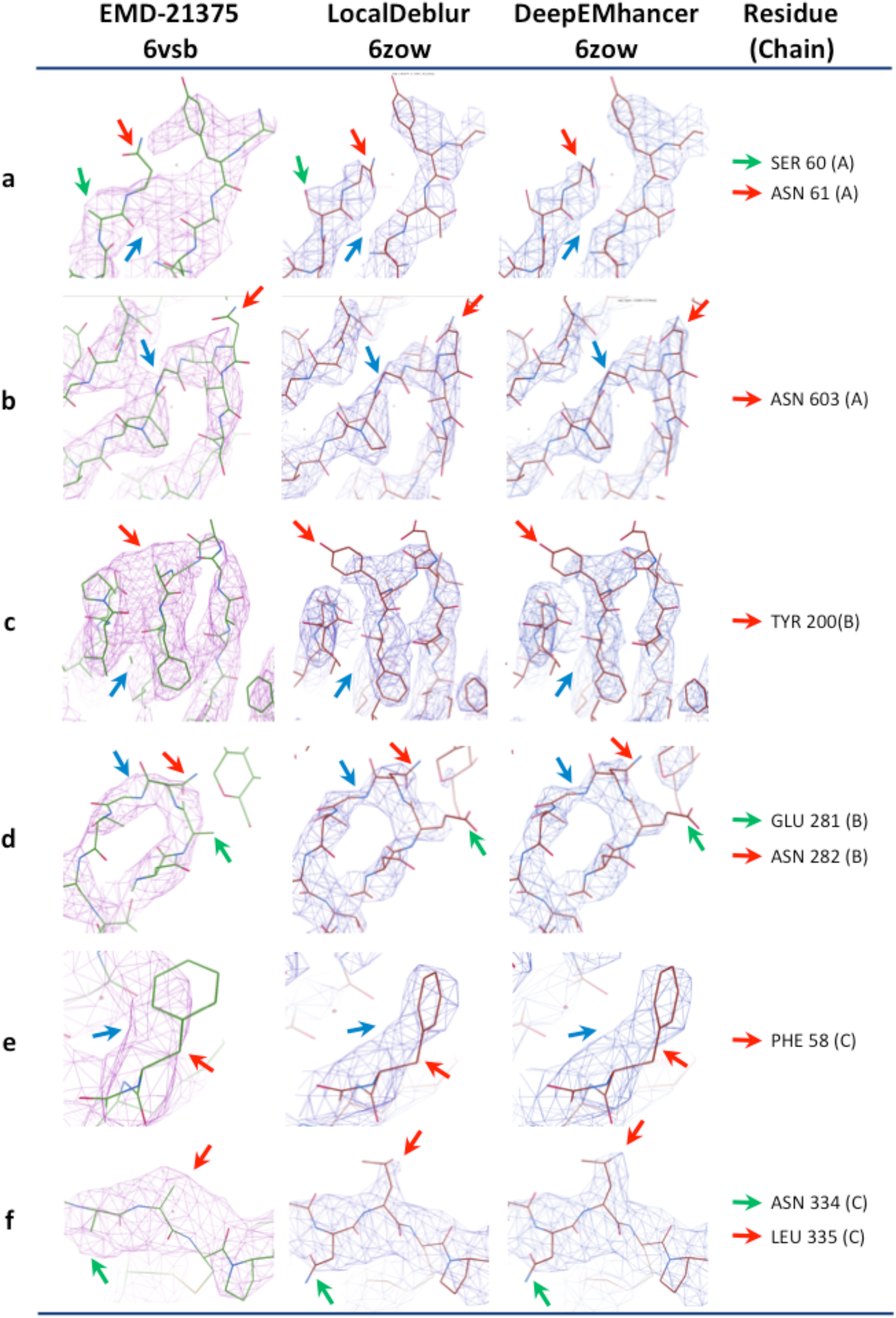
Comparison of the ability to trace the atomic structure between the original cryo-EM map (EMD-21375) and the two sharpened maps derived from the new reconstructed ensemble map. Six representative 3D map areas (a-f) illustrate the fitting between map and atomic structure. The red arrows detail aminoacid side chains fitted to better defined densities in the sharpened map compared to the original map. These side chains could have been modeled (a, b, d, e) or being absent (c, f) in the original map. The green arrows indicate other additional residues whose side chains have been modeled only in the sharpened maps, while they were absent in the original one. The blue arrows point at densities that make it difficult to follow the carbon skeleton shape or discriminate among different chains in the original map, whereas they appear better resolved in the sharpened maps.

**Supplementary Material Figure SM3.**
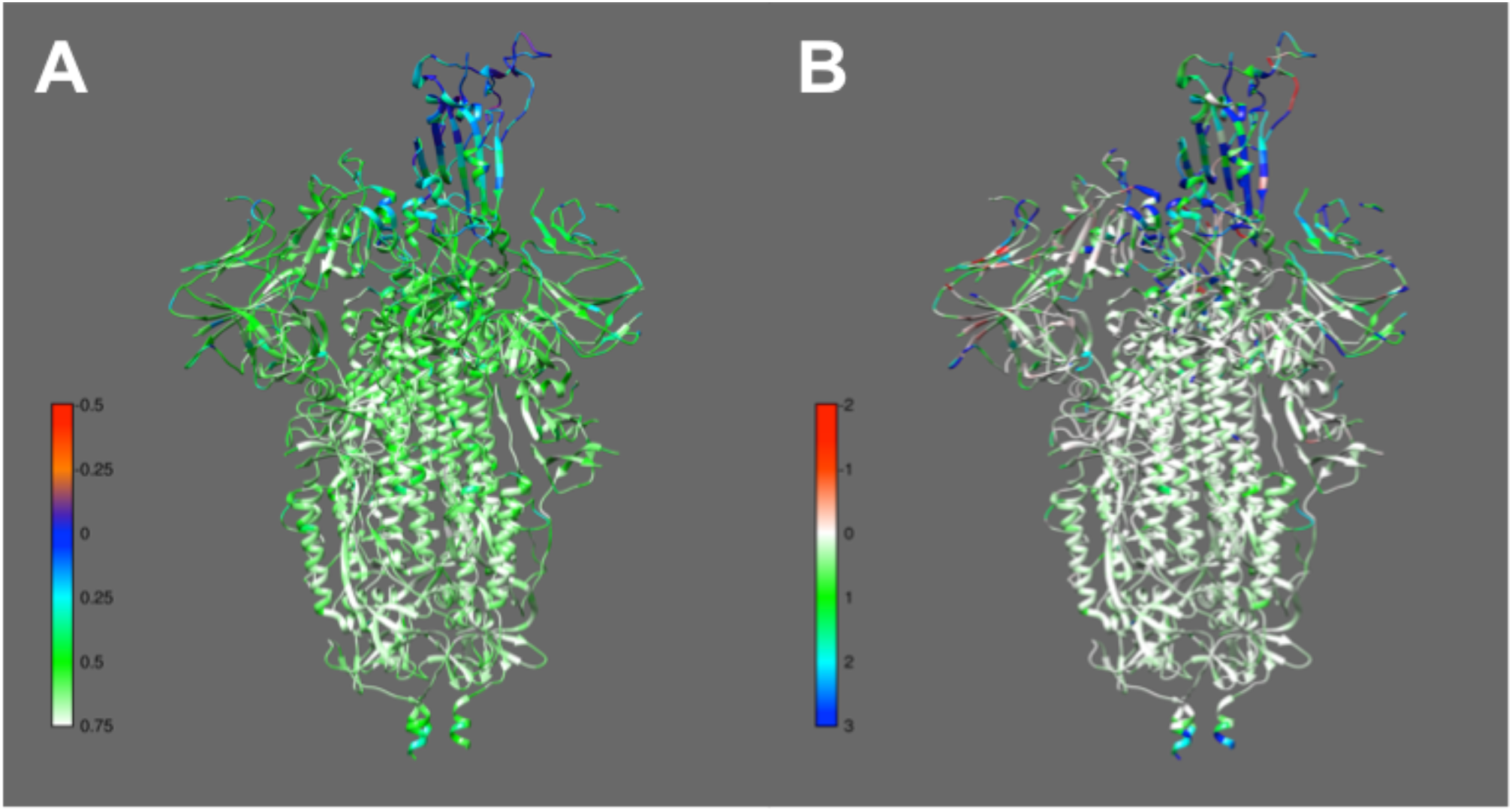
Map-to-Model quality measures for the ensemble map. A) Q-score values represented on each amino acid of the new ensemble atomic model. Q-scores close to 1 indicate better resolvability of the residue. B) FSC-Q values are represented for each residue. Values close to zero indicate a good map-to-model fit, while values far from zero indicate areas where the model loses support with respect to the map signal.

**Supplementary Material Figure SM4.**
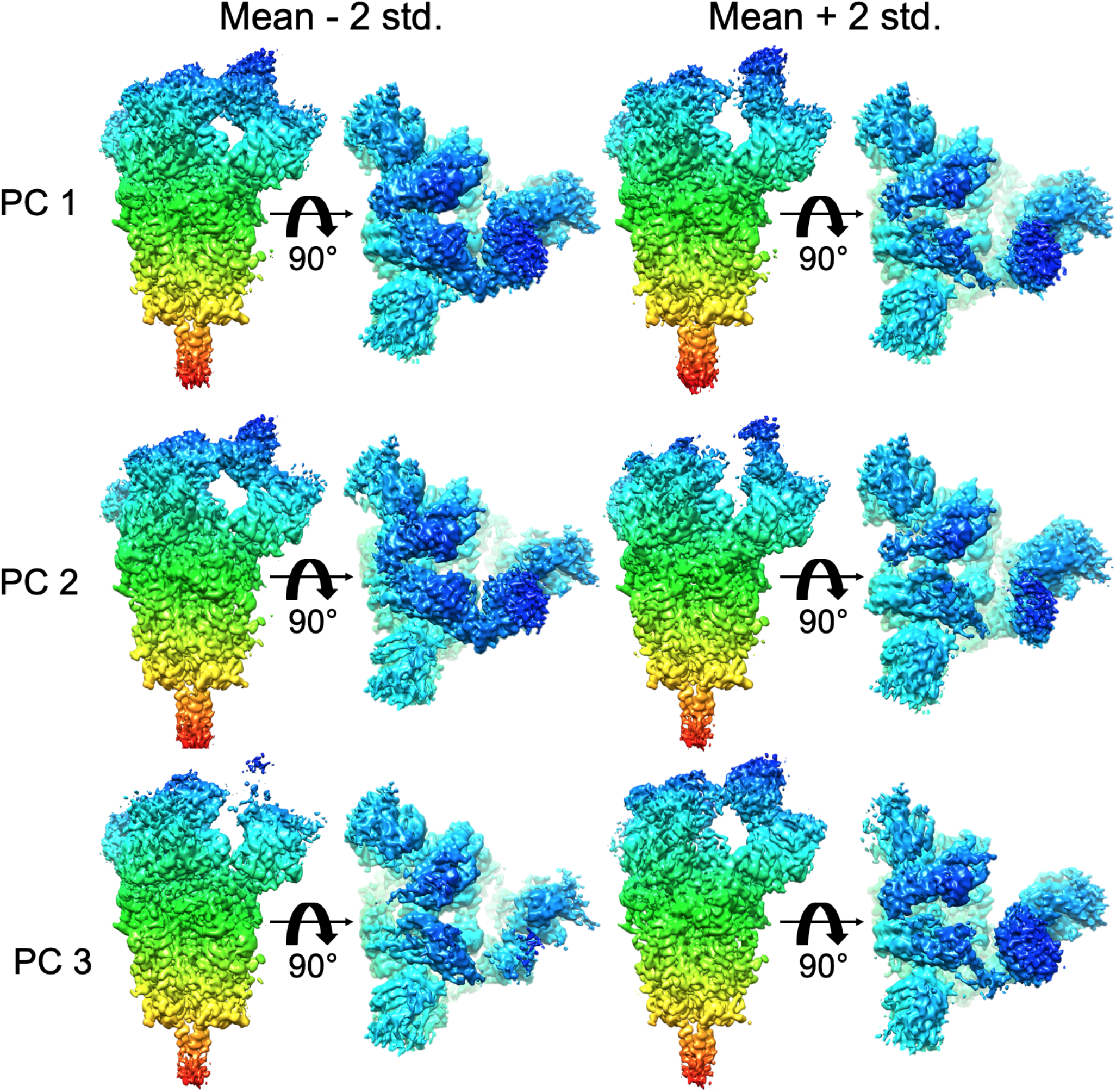
Principal Component Analysis. Side and top views of mean volume +/-2 std for the three principal components. Coloring indicates z-depth of the structure, and is added to assist visualization of the top view.

**Supplementary Material Figure SM5.**
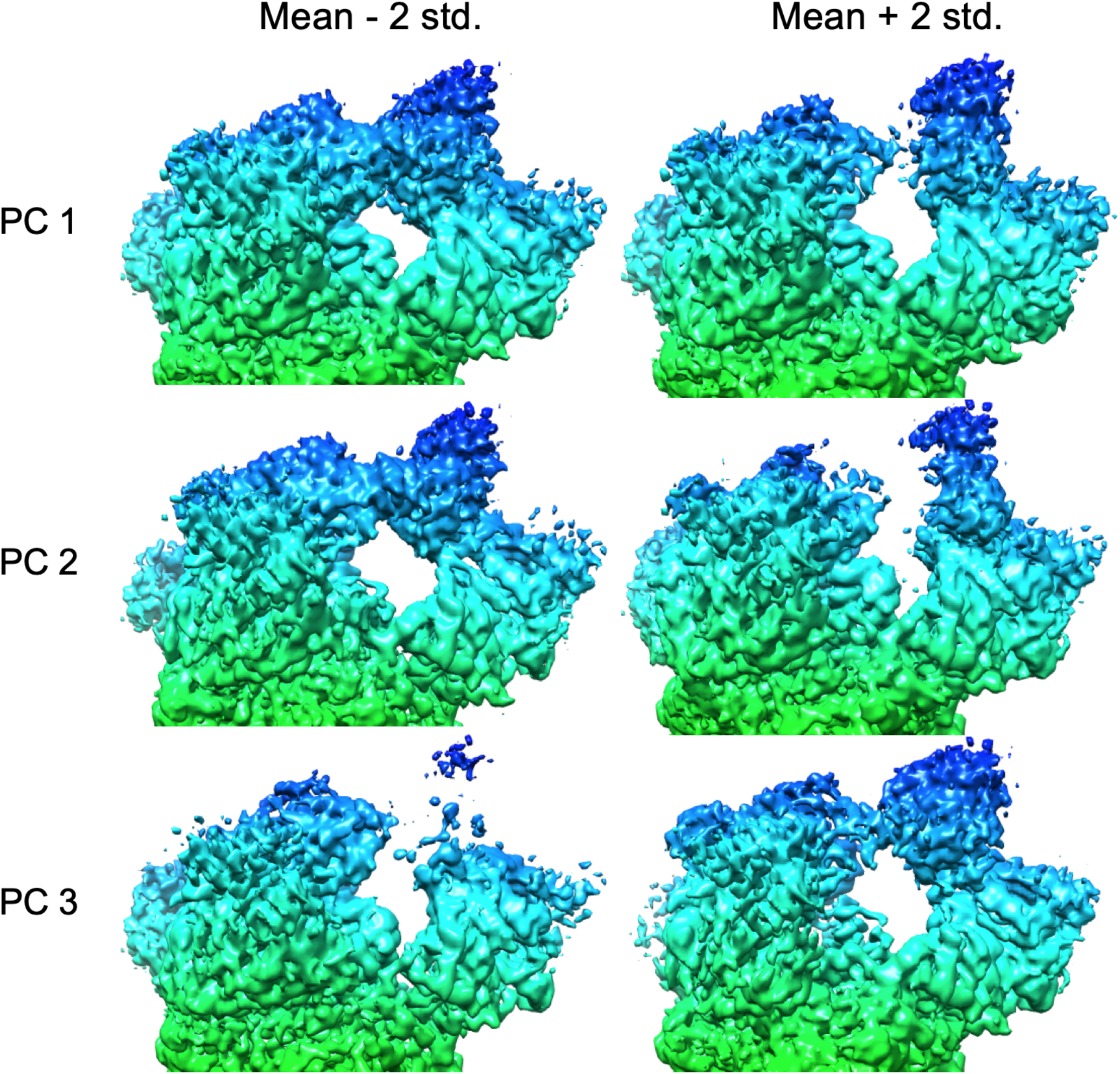
Principal Component Analysis. Enlarged top half of the side views showing details of the density changes captured by the principal components. Coloring indicates z-depth of the structure, and is added to assist visualization of the top view.

**Supplementary Table SM1.** Validation scores of the new atomic structure PDB 6ZOW

| Validation scores | Whole structure<sup>1</sup> | Modeled structure<sup>2</sup> |
| --- | --- | --- |
| EMRinger | 2.66 | 2.73 |
| CC(mask) | 0.75 | 0.76 |
| Ramachandran outliers (Goal: < 0.2%) | 0.00 | 0.00 |
| Ramachandran favored (Goal: > 98%) | 97.06 | 97.28 |
| Rotamer outliers (Goal: < 1%) | 0.52 | 0.17 |
| C-beta outliers | 0 | 0 |
| Clashscore | 12.06 | 12.43 |
| MolProbity overall score | 1.76 | 1.74 |
| Q-Score | 0.55 | 0.58 |
| FSC-Q | 0.73 | 0.58 |

<sup>1</sup> Chains A, a, B, C
<sup>2</sup> Excluding the chain a (RBD, added by rigid fitting)

**Supplementary Table SM2.** Number of sequons and size of their respective glycan chains.

| Subunit | Size of N-linked glycan chain | 6VSB |  | 6ZOW |  |
| --- | --- | --- | --- | --- | --- |
|  |  | Number of sequons | Number of glycans | Number of sequons | Number of glycans |
| S1 | 1 | 26 | 26 | 18 | 18 |
|  | 2 | 0 | 0 | 7 | 14 |
|  | 3 | 0 | 0 | 2 | 6 |
|  | **Subtotal** | **26** | **26** | **27** | **38** |
| S2 | 1 | 1 | 1 | 0 | 0 |
|  | 2 | 17 | 34 | 9 | 18 |
|  | 3 | 0 | 0 | 5 | 15 |
|  | 4 | 0 | 0 | 2 | 8 |
|  | 5 | 0 | 0 | 2 | 10 |
|  | **Subtotal** | **18** | **35** | **18** | **51** |
| **TOTAL** |  | **44** | **61** | **45** | **89** |

**Supplementary Material Movie 1**. Movie presenting the morphing between the two algorithmically stable classes described in the main text, spanning Principal Component Axis 1.

## Notes

### Competing Interest Statement

The authors have declared no competing interest.

